# Evolutionary divergence of anaphase spindle mechanics in nematode embryos constrained by antagonistic pulling and viscous forces

**DOI:** 10.1101/2021.08.10.455863

**Authors:** Dhruv Khatri, Thibault Brugière, Chaitanya A. Athale, Marie Delattre

**Affiliations:** Div. of Biology, IISER Pune, Dr. Homi Bhabha Road, Pashan, Pune 411008, India; Laboratory of Biology and Modeling of the Cell, Ecole Normale Supérieure de Lyon, CNRS, Inserm, UCBL, 69007 Lyon, France

**Keywords:** evolutionary cell biology, cell mechanics, forces, nematodes, cytoplasmic viscosity, cryptic evolution, spindle positioning, viscoelasticity

## Abstract

Cells are the basic unit of biological organization, and their division is remarkably conserved across phyla. However from an evolutionary perspective, it remains unclear how much cellular parameters can diverge, without altering the basic function they sustain. We address the mechanics of asymmetric mitotic spindle positioning during the first embryonic division of six nematode species. We propose a viscoelastic model of spindle positioning and mobility that can provide a physical explanation of why in *C. elegans* it undergoes oscillations during elongation, whereas most others lack oscillations. To test this model, we measured the pulling forces and opposing cytoplasmic drag by a combination of laser ablation of the anaphase spindle and tracking of intracellular granules. While centrosomes of all species recoil on spindle cutting, quantitative differences correlate with the cytoplasmic viscosity. In fact, increased viscosity correlates with decreased oscillation speeds of intact spindles across species. However, the absence of oscillations despite low viscosity in some species, can only be explained by smaller pulling forces. Consequently, we find that spindle mobility across the species analyzed here is characterized by a tradeoff between cytoplasmic viscosity and pulling forces. Our work provides a framework for understanding mechanical constraints on evolutionary diversification of spindle mobility.

## Introduction

The ubiquitous nature of physical principles means it is expected that they influence cell physiology as seen in the role of mechanics of the cytoskeleton-motor systems in cell division, transport and regulation and influence of diffusion. This would also suggest these physical principles of diffusion, mobility and transport must also constrain the evolutionary diversification of these processes. For example the size of spindles has been seen to linearly scale with cell size during embryogenesis across metazoan species (Crowder et al., 2015), but the biophysical mechanism that governs such apparently universal scaling has proven harder to identify, potentially due to the limitations of physical quantification in species beyond a relatively narrow set of model species. Additionally the variation in measured properties could result from drift with no adaptive significance, as seen in genomes, proteomes and network evolution (Lynch, 2007). As a result, genetic variation might accumulate without any effect on phenotypic property. Therefore examining to which extent physical properties can evolve without constraints on phenotypic variation, remains to be explored.

In an attempt to address this, we had previously characterized the first embryonic cell division in 40 closely related species of nematodes (Valfort et al., 2018). In all species, the division is asymmetric due to the asymmetric positioning of the mitotic spindle towards the posterior side of the cell. However, the movements of the spindle during its displacement are very different from one species to the other, suggesting cryptic changes in the cellular parameters that govern spindle motion.

In nematodes, the anterior/posterior (A/P) polarity of the animal is established after fertilization. One manifestation of this symmetry breaking is the asymmetric displacement of the mitotic spindle during the first anaphase, from a central to a posterior position. Consequently, the division generates two daughter cells of unequal size and of unequal fate. In the nematode *C. elegans*, during this displacement, the spindle also undergoes vigorous movements that are perpendicular to the A/P axis. Centrosomes oscillate back-and-forth along the transverse axis at a specific frequency and amplitude, while the anterior and the posterior centrosomes move in a manner that mimics anti-phase oscillations (offset by half a wavelength). These stereotypical movements are referred to as spindle oscillations. Laser ablation of the central spindle at the onset of anaphase in *C. elegans*, resulted in both centrosomes accelerating towards the cell poles, demonstrating that opposite pulling forces act on astral microtubules (MTs) to displace the centrosomes (Grill 2001). These movements result from the activity of a conserved dynein-containing protein complex anchored at the cortex (Kotak, 2019). Inactivation of this complex greatly affects spindle elongation, spindle displacement and spindle oscillations, suggesting that cortical pulling forces control all aspects of spindle motion in *C. elegans* embryo. While, an incomplete activation of this complex does not affect the asymmetric cell division, it abolishes spindle oscillations. It has thus been proposed that oscillations emerge above a threshold of active forces (Pecreaux et al., 2006). Oscillations have also been quantitatively reproduced by physical models and simulations (Grill et al., 2003, Grill et al., 2005, Kozlowski et al. 2007). Cortical force generators (dynein-containing complexes) pull from each side of the cortex (upper and lower), which should leave the centrosome in a stable, central position. In both models, a positive feedback mechanism is then implemented to recapitulate the transverse displacement of the centrosomes. A slight displacement of the centrosome towards the upper cortex for instance, is amplified because pulling forces increase as the centrosome comes closer to the cortex. This could happen for instance if the load per motor decreases as the distance to the centrosome decreases (Pecreaux et al., 2006). Next, the centrosome will go back to the center of the cell because of a restoring force. This force could be generated by astral MTs pushing on the cortex as they polymerize (Kozlowski et al, 2007), or by the buckling of these MTs, extending laterally to the oscillation axis (Pecreaux et al., 2006). The “tug-of-war” between these pulling and restoring forces generates the oscillations. Pulling forces must also counter-balance the damping force generated by the viscous cytoplasm in order to launch the oscillations. Spindle motion is thus caused by the complex interplay between different mechanical forces and material properties of the cell and of the spindle.

In previous work, we recorded DIC microscopy time series of 40 nematode species belonging to the *Caenorhabditis* genus, or to closely related genera (Valfort 2018). Although they all undergo an asymmetric first cell division, some species show clear quantitative differences in spindle motion when compared to *C. elegans*. Interestingly, spindle transverse oscillations are restricted to *Caenorhabditis* species. Only in *C. monodelphis* which is the most basal *Caenorhabditis* species, and in all species outside of this genus, the anaphase spindle is asymmetrically displaced without any transverse oscillations. Here, we asked which cellular parameter change accounts for this absence of oscillations. A simple hypothesis is that the viscosity of the cytoplasm could give rise to these differences. Indeed, predictions from simulations of *C. elegans* spindle-oscillatory mechanics have suggested order of magnitude differences in the cytoplasmic viscosity can change the qualitative nature of spindle oscillations by mechanical damping (Kozlowski et al., 2007). However, these predictions remain to be tested experimentally, since altering cellular viscosity without affecting cell physiology is technically challenging. Alternatively, if the net pulling forces are reduced, we also expect the loss of oscillations. Across species, spindle oscillation buildup may be hindered by an increase in cytoplasmic viscosity or by a reduction in cortical forces arising from gene expression changes or reduced cortical localization or other parameter change. Multiple such scenarios of evolutionary change can be envisioned, and with each species having its own combination of changes.

In this study, we address the question of how many biophysical features explain the diversity of spindle motion over the course of nematode evolution. We chose 6 representative species to specifically ask how variable the cytoplasmic viscosity between species is. We also independently measure the cortical pulling forces responsible for spindle motion in all species. We examine which of the changes correlate with the observed difference in spindle motion.

## Results

### Species dependent variation in spindle motion of related nematode embryos during the first asymmetric cell division

We chose to explore which cellular parameters can explain the absence of spindle transverse oscillations in some nematode species. Among those previously described (Valfort & al. 2018), we selected 4 species without spindle oscillations belonging to 4 distinct genera: *Pristionchus pacificus, Oscheius tipulae, Diploscapter species 1 JU359 (D. sp. 1)* and *Caenorhabditis monodelphis*. We also chose *C. remanei and C. elegans* as control species displaying anaphase spindle oscillations (Fig. 1). These species are also characterized by variations in cell size. For instance, *P. pacificus embryos* are 10% longer and *D. sp. 1* are 20% shorter than *C. elegans* embryos (Table S1). As previously shown on a larger set of species, the lack of spindle oscillation is not restricted to small or large embryos, thus cell size change alone is unlikely to be responsible for evolutionary changes in spindle oscillations (Valfort et al., 2018). We also estimated cell cycle length by measuring the time spent between the first nuclear envelope breakdown and the onset of the first cytokinesis, from the time-lapse recordings. We found the cell cycle was the shortest in *C. elegans* and *C. remanei. O. tipulae* and *P. pacificus* are 1.7 slower than *C. elegans*, whereas *C. monodelphis* and *Diploscapter sp*. 1 are 2.2 and 3 times slower than *C. elegans*, respectively (Table S1), raising the possibility that spindle oscillations are restricted to rapidly dividing species.

**Figure 1.**
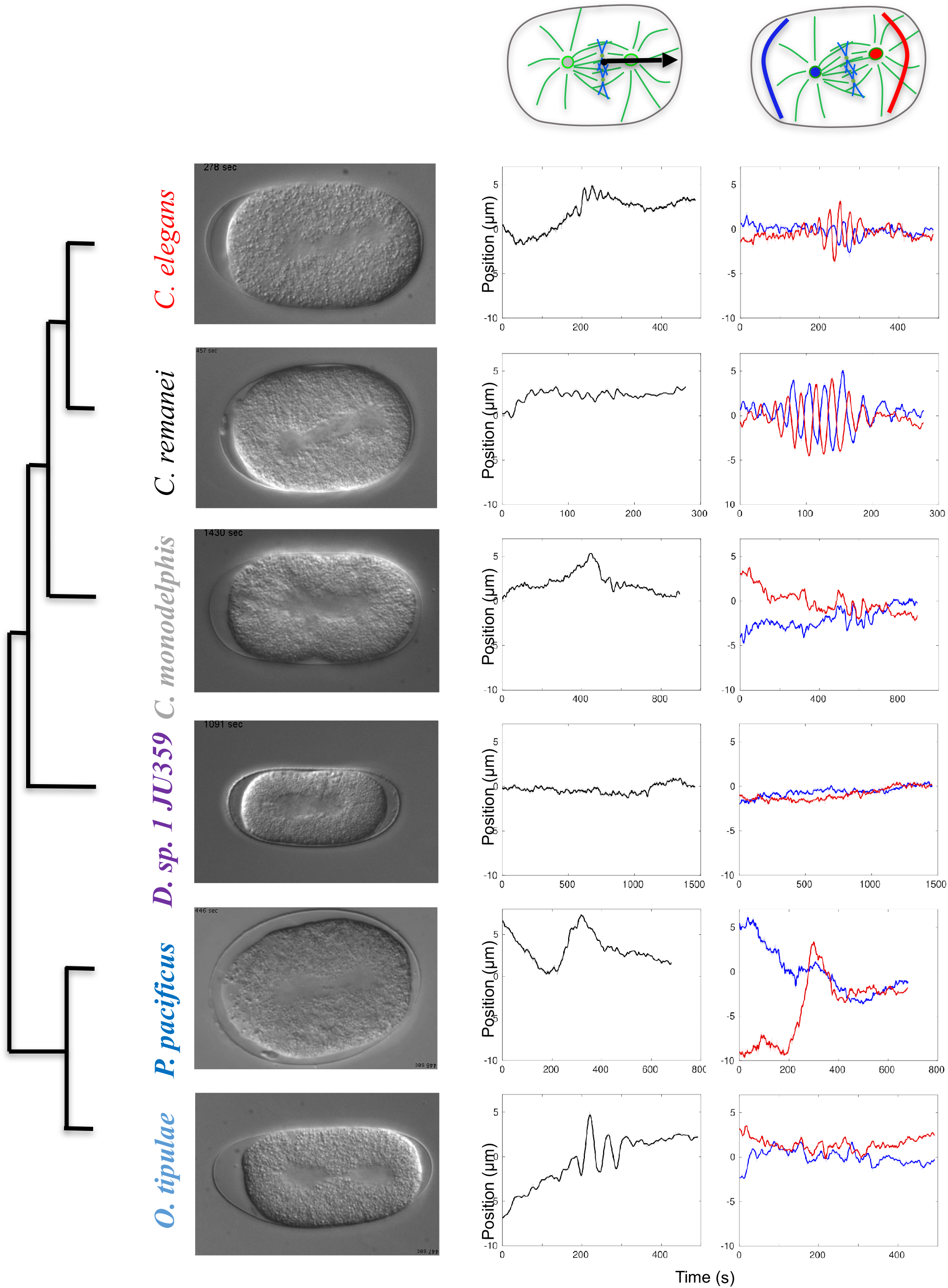
Diversity of spindle dynamics across nematodes. The diversity in spindle movement in single celled embryos of related nematode species is seen in the DIC images. These images were used to estimate the change in position of the spindle mid-plane along the AP axis (black lines) and the transverse movements of the spindle poles (blue: anterior, red: posterior). Some of the species display oscillations of spindle poles in the transverse direction, while some even display AP oscillations.

### Dynamics of anaphase spindle pulling varies independently of oscillations across species

Previous work has shown spindle movement, including oscillations, is mainly driven by pulling forces acting on the spindle in *C. elegans* embryos (Grill et al., 2001). However, in some distant species, for instance in the yeast S. pombe (Tolic et al., 2004) and during meiotic divisions in some cell types (for instance oocytes meiotic divisions in most species), spindle positioning is independent of microtubule-based pulling forces from the cortex (Almonacid et al., 2014). We first asked whether the absence of spindle oscillations in some nematode species reflects a mechanism that is independent of pulling forces. Mechanical forces acting on the spindle can be revealed by laser ablation of the central spindle at the onset of mitosis. Following spindle severing, the centrosomes recoil towards the cell pole if they are initially pulled (Grill et al., 2001). This is because the central spindle connects the poles and holds the balance, much like a stretched rubber-band. In species in which the mitotic spindle elongates by inside-out pushing forces, spindle severing leads to the collapse of the centrosomes at the center of the spindle (Khodjakov et al., 2004, Tolic et al., 2004).

We used a pulsed UV laser to sever the spindle at the onset of anaphase in all the 6 species (Fig. 2A, Fig. S1) and analyzed the recoil trajectories of the anterior and the posterior centrosomes after the cut. The centrosomes of most of the species appear to recoil towards the cell pole (Fig. 2B). However the movement of *D. sp. 1* centrosomes was very limited with the characteristic recoil time being almost as long as the time of acquisition (27 s) with τ = 23.7 and 23 seconds, anterior and posterior respectively (Fig. 2C) and velocity of ∼ 0.05 and 0.07 μm/s, anterior and posterior poles (Fig. 2D). The half time of recoil is ∼5 s for *C. elegans* and *C. remanei*, while the remaining 4 species have longer times (Fig. 2B). The recoil velocity of centrosome translocation in *C. elegans* is ∼1.1 μm/s (anterior) and ∼1.2 μm/s (posterior) (Fig. 2D, Table S2), comparable to previous reports (Grill et al., 2001). From these results we concluded that the mitotic spindle is subjected to cortical pulling forces during anaphase in all species.

**Figure 2.**
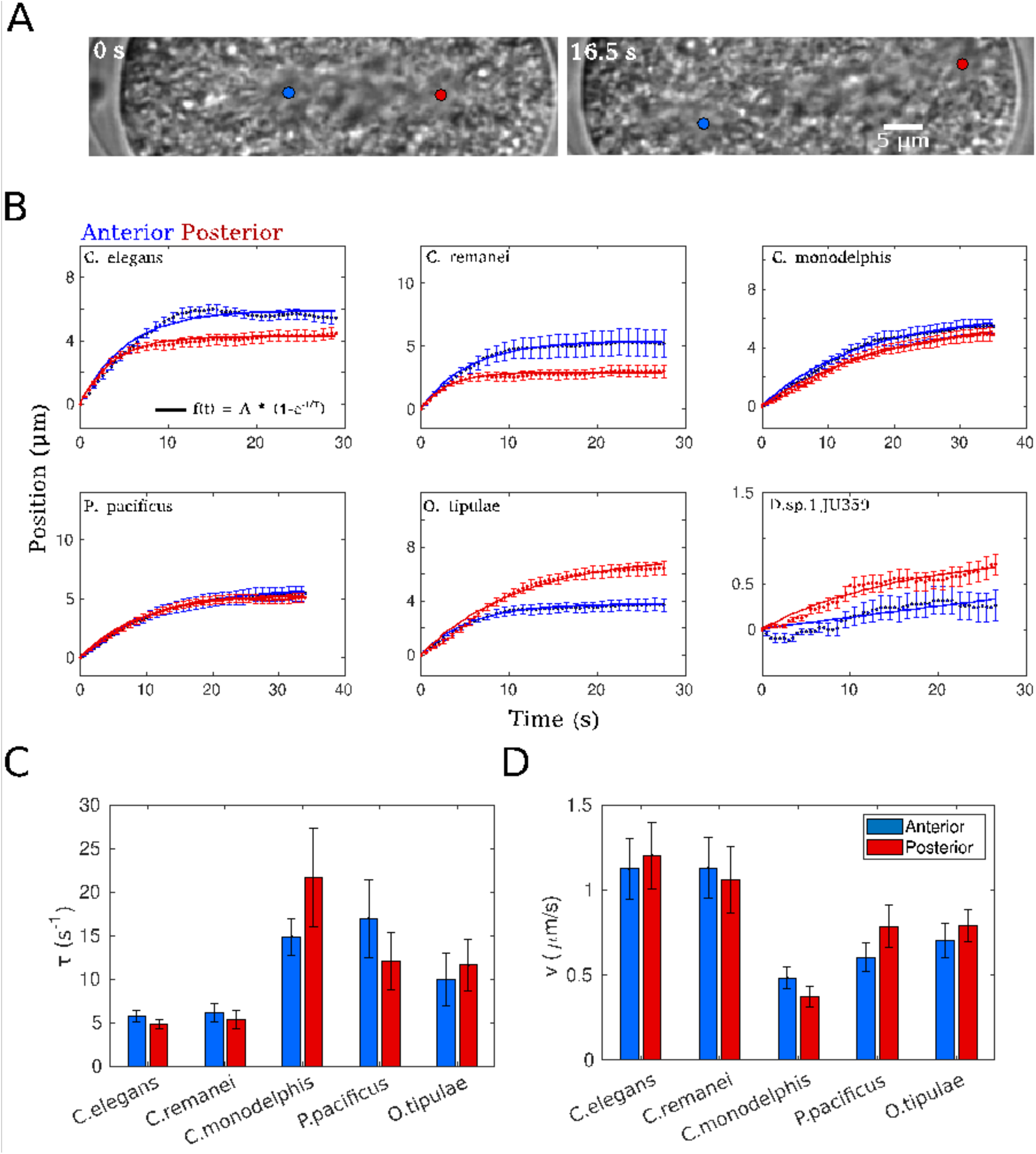
Centrosome trajectories post laser ablation characterised by recoil velocity and decay constant. **(A)** Representative images of a *C. elegans* embryo before (0 s) and 16.5 s after laser of the spindle, with the anterior (blue circle) and posterior (red circle) centrosmes marked. **(B)** Recoil trajectories of the anterior (blue) and posterior (red) centrosomes after laser ablation for different species were averaged (filled circles) with error bars indicating SEM. The data was fit to the recoil model (bold line) given by Equation 8. The individual profiles are also plotted (thin lines). **(C**,**D)** The fit parameters for each species in terms of **(C)** the decay constant τ (1/s) and **(D) recoil** velocity v (μm/s) are plotted (mean± sem). Colours indicate anterior (blue) and posterior (red) trajectories.

Nevertheless, the recoil trajectories of centrosomes after the cut vary between species in terms of the recoil velocity, half-time of recoil, final position of the centrosome and anterior and posterior differences (Fig. 2B, C). For all species that do not display oscillations, the half time of recoil was lower than those of *C. elegans*, whereas the initial velocity was systematically lower.

### Cytoplasmic viscosity can change by an order of magnitude between closely related species

Visual inspection of the DIC image time-series suggested qualitative correlation between spindle mobility patterns and passive mobility of these granules. Additionally, from first principles of fluid mechanics, the mobility of intracellular organelles and structures is expected to experience viscous drag and be an important determinant in their motion. The spindle mobility differences could thus most simply be explained by evolutionary changes in cytoplasmic viscosity, and provide a direct link to spindle motion. To test this hypothesis we proceeded to estimate the cytoplasmic viscosity across the species.

In DIC images, nematode embryo cytoplasm is prominently packed with clearly visible yolk granules (Clokey and Jacobson, 1986; Hermann et al., 2005). We decided to estimate cytoplasmic viscosity in the different species using granule mobility, as previously used for *C. elegans* embryo (Grill et al., 2001). Visual inspection of the DIC image time-series of the embryos suggested qualitative correlation between spindle mobility patterns and passive mobility of these granules. Since granules undergo streaming due to spindle movements and are at times even actively transported, we followed granule mobility during interphase and in the top plane of the embryos far from the spindle plane. This was done to minimize the effects of active transport on the granule mobility measurement, so we expect it to be largely diffusive, i.e. thermal random motion (Fig. 3A, see Material and Methods). Embryo images were partitioned into anterior, middle and posterior regions and over 1,000 granules in the anterior posterior portion of each embryo were tracked (Fig. 3B) using a previously developed MATLAB code for single particle tracking in DIC images (Chaphalkar et al., 2021). Multiple embryos of each species were analyzed and the mean square displacement (MSD) of granules calculated (see Material and Methods, Equation 4). Granule MSD plots were averaged over time and across multiple granules and fit to the anomalous diffusion model (see Material and Methods, Equation 5). The anomalous diffusion model includes a MSD ∼ t^α^ dependence, which if α ∼ 1 results in a linear dependence indicating of normal diffusion, the slope of the line indicates the diffusion coefficient (Athale et al., 2014; Khetan and Athale, 2016). Based on the fit value of 0.96 of the anomaly parameter, we conclude that granule movement was effectively diffusive in both anterior and posterior regions as well as the related species (Fig. S2). We estimate the radius of lipid granules from each species averaged across multiple individuals (Fig. S3) and determine the solvent phase viscosity (η_s_) using the Stokes-Einstein relation, see Material and Methods, Equation 6 (Einstein, 1905; Berg, 1993). The granules themselves appear to be highly packed in the cytoplasm and the packing appears to differ between species. Crowding of the cytoplasm affects the effective viscosity. To account for it we measured an area packing ratio (ϕ_2D_) of the granules in the anterior and posterior regions based on their number and size (Fig. 3D and Methods section) and substituted it in Equation 7 (see Material and Methods) to estimate the effective viscosity (η_eff_) that accounts for self-crowding effects using soft-sphere packing theory (Quemada, 1977). Using this approach, we found a cytoplasmic viscosity for *C. elegans* of 0.67 Pa s averaged for both halves of the embryo, in the same range as previously reported values ranging from 0.1 (Garzon-Coral et al. 2016) to 1 Pa s (Daniels et al., 2006).

**Figure 3.**
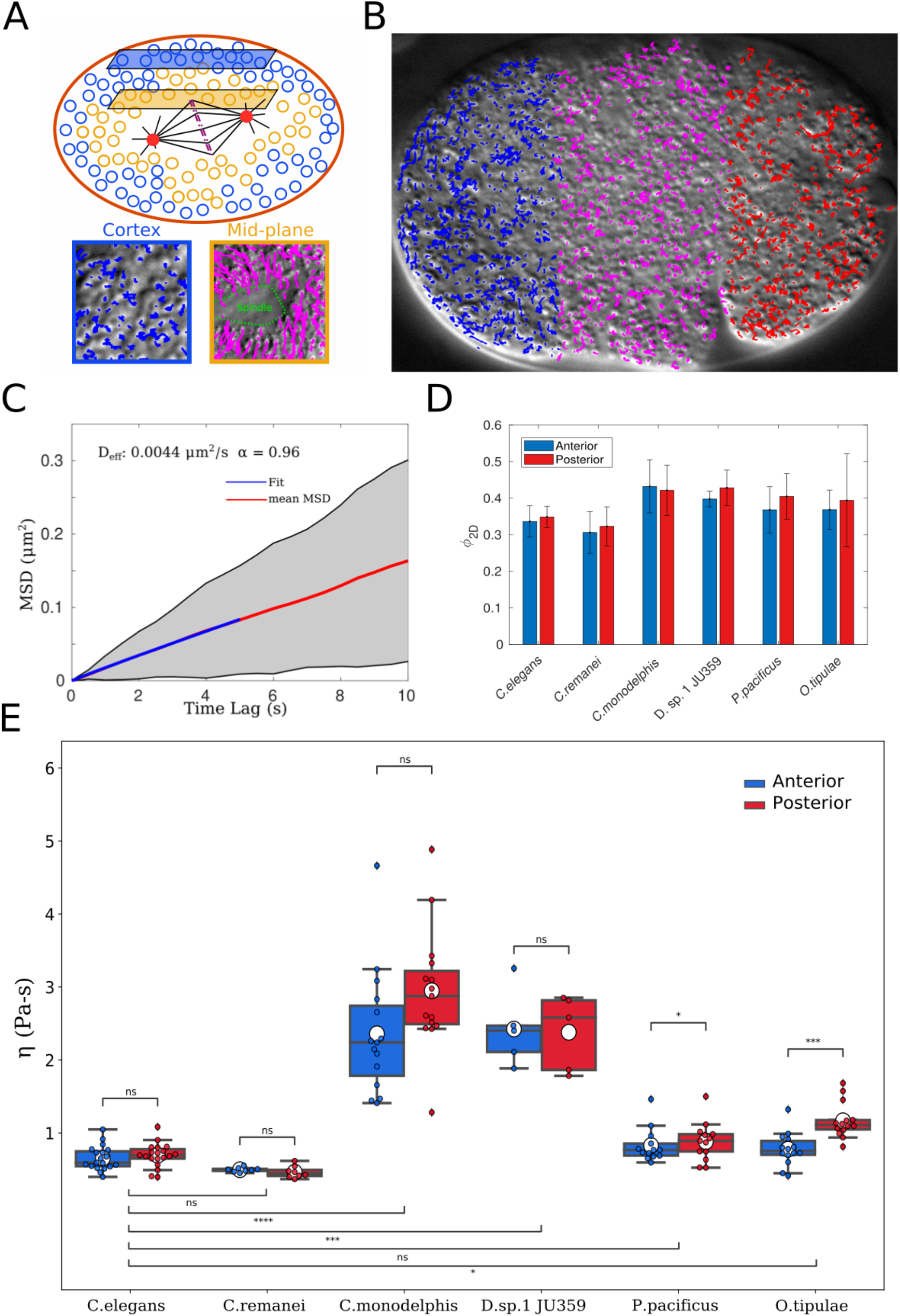
Cytoplasmic viscosity estimates from yolk granule mobility. **(A)** Schematic of metaphase spindle and granules present in the cytoplasm of a one-celled nematode embryo. Granules can be distinguished in terms of their motility based on their position in the embryo-those below the cell cortex (blue) and those in close proximity to the spindle (yellow). Two different planes of DIC image acquisition were then used to track the granules, and representative results from the cortex (blue) and mid-plane (yellow) suggest a qualitative difference in motility. **(B)** The granules of the cortical region of a *C. elegans* embryo in DIC were tracked from time-series and the tracks were classified into anterior (blue), middle (magenta) and posterior (red) regions by considering 1/3 of the major axis. **(C)** These trajectories were used to generate mean square displacement (MSD) curves for the whole embryo as a function of time. The average MSD (red) line was fit to the anomalous diffusion model fit (blue, Equation 4) and fit parameters of D_eff_ = 0.0044 μm^2^/s and α = 0.96 obtained. **(D)** The mean±SD area packing fraction of granules (ϕ_2D_) from the anterior (blue) and posterior regions (red) is plotted for the 6 nematode species analyzed. **(E)** The effective viscosity η_eff_ of each nematode species from the anterior (blue) and posterior (red) regions are plotted with mean (white circle), median (horizontal line) and 1st and 2nd quartiles (whiskers) indicated. Points indicate outliers. The KS test of significance of inter-species and anterior/posterior differences was applied. The asterisks indicate * p<0.05, ** p<0.01, *** p<0.001 and **** p<0.0001 while ns: not significant.

The viscosity estimation is based on the validity of diffusion as the primary process driving granule motion, i.e. passive, thermal Brownian motion. If active intracellular movements also affect the mobility blocking intracellular ATP production should result in a change in the measure of mobility. *C. elegans* embryos were treated with RNAi against *atp-2* (ATP synthase subunit) or *cyc-1* (Cytochrome-c1), shown previously to inhibit mitochondrial ATP production (Tsang et al, 2001; Neves et al., 2015). Compared with untreated *C. elegans* embryos, we found no quantitative difference in granule mobility (Fig. S4A), MSD profiles (Fig. S4B), estimated diffusion coefficient or viscosity (Fig. S4C,D). This strongly supports the hypothesis that granule mobility we report here is indeed diffusive.

The cytoplasmic effective viscosity (η_eff_) differed between species. Except for *C. remanei*, all species showed viscosity values higher than *C. elegans*, with *C. monodelphis* showing the highest viscosity of 2.65 Pa s (Fig. 3E, Table 1). Nevertheless, the difference between *C. elegans* and *P. pacificus* was not statistically significant, most likely due to high inter-individual variability in *P. pacificus*. We also find viscosity differs greatly between anterior and posterior regions in *O. tipulae*, and to a lesser extent in *P. pacificus*. The viscosity in the anterior and posterior halves of the embryos were however similar in *C. elegans, C. remanei, C. monodelphis* and *D. species 1* (Fig. 3E and Table 1). These results reveal even closely related species show a diversity in cytoplasmic viscosity with an almost 10-fold difference between the most extreme values measured (i.e. one order of magnitude).

**Table 1:** Quantification in the divergence between nematode embryos with regard to cell size, centrosome size and the duration of the cell cycle.

| Table 1 |  |  |
| --- | --- | --- |
| *Species* | *Effective Viscosity (Pa s)* |  |
|  | *Anterior* | *Posterior* |
| *C. elegans* (n = 19) | 0.654 ± 0.165 | 0.694 ± 0.162 |
| *C. remanei* (n = 8) | 0.496 ± 0.031 | 0.464 ± 0.083 |
| *C. monodelphis* (n = 15) | 2.359 ± 0.858 | 2.947 ± 0.833 |
| *D. sp. 1, JU359* (n = 5) | 2.424 ± 0.521 | 2.379 ± 0.520 |
| *P. pacificus* (n = 13) | 0.828 ± 0.233 | 0.881 ± 0.263 |
| *O. tipulae* (n = 14) | 0.778 ± 0.229 | 1.168 ± 0.242 |

While the increase in viscosity across species (Fig. 3E) qualitatively correlates with slower recoil dynamics in spindle cutting cutting experiments (Fig. 2C, D), it is unclear if that can explain the absence of bonafide spindle oscillations in some species such as *P. pacificus*, since the cytoplasmic viscosity measured is statistically similar to that of *C. elegans*. Therefore we proceeded to examine whether the magnitude of the pulling forces on the spindle may also vary between species.

### The net pulling forces and elasticity forces acting on centrosomes also vary between closely related nematode species

While spindle laser ablation experiments have been used in the past to estimate relative rates of pulling by forces acting on the astral MTs, the quantification of absolute forces requires a mechanical model of the spindles, motors and the cytoplasm. In recent work spindle centering in metaphase of *C. elegans* embryos has modeled to infer restoring forces on the centrosomes by considering both elastic and viscous components, i.e. a viscoelastic model of movement (Garzon-Coral et al. 2016). Indeed multiple studies point to cytoplasmic mechanical properties to be best explained by viscoelasticity (Berret et al. 2016, Fabry et al., 2001). The Kelvin-Voigt (KV) model is invoked to account for not just the spring force (F) that acts on laser ablated centrosomes pulling them backwards, but also the viscous drag (γ) due to the presence of cytoplasm, crowding by granules and the elasticity (k) of the half-spindle and actin meshwork (Fig. 4A). The fact that most of the 6 species tested show a recoil of centrosomes on ablation, suggests we can use the same physical model to understand the mechanics across species. Based on the KV-model the position of the centrosome as function of time, p(t) is fit to the following equation:

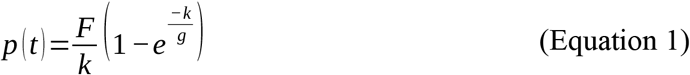

**Figure 4.**
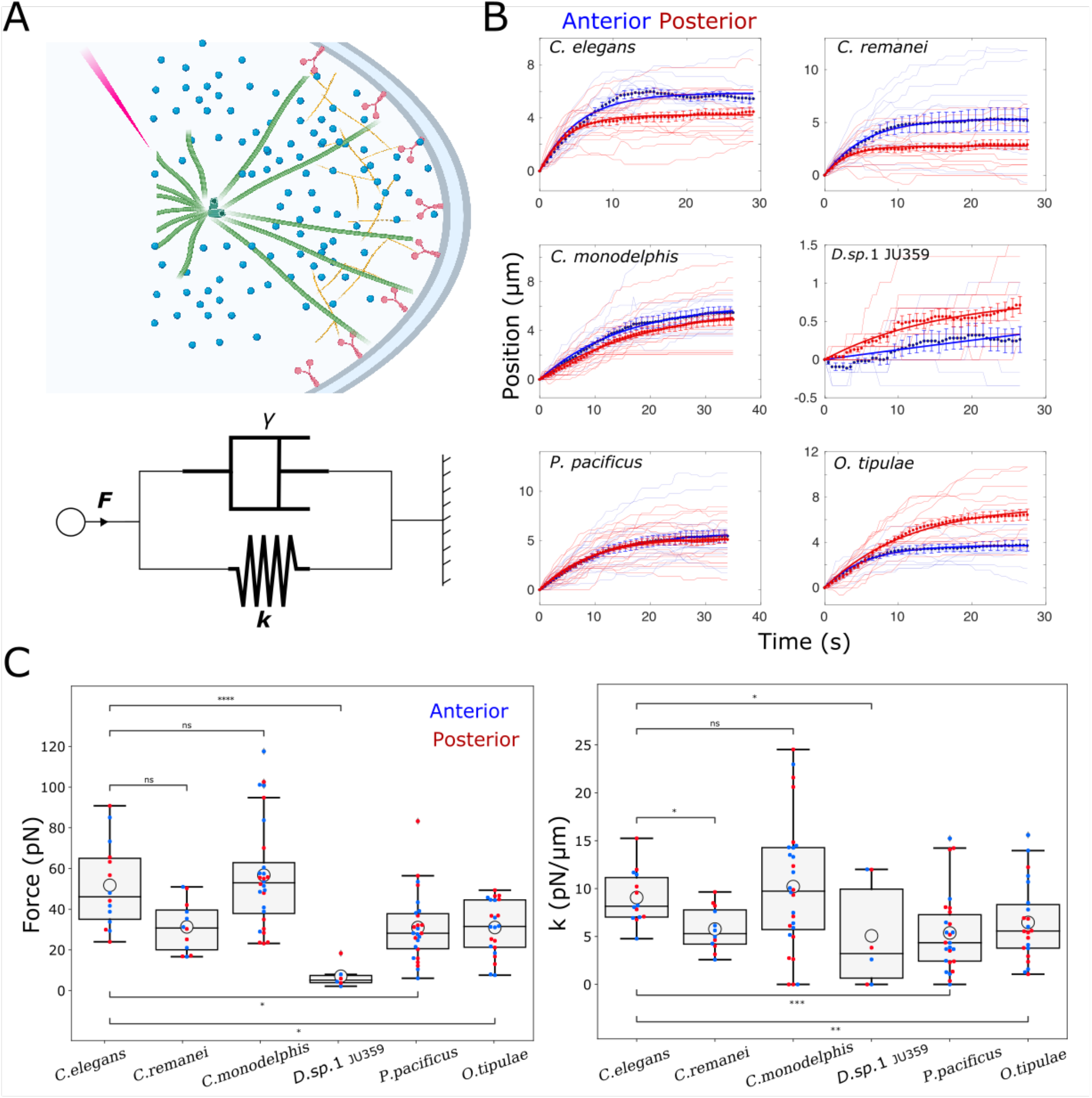
Pulling forces estimated from viscoelastic model fit to recoil data. **(A)** Schematic of laser ablation of the spindle mid-plane that results in recoil motion of the centrosome modeled by the Kelvin-Voigt viscoelastic model represented here by a spring with elasticity k and dashpot with viscous drag coefficient γ, that acts to damp movement due to the pulling force, F. **(B)** Recoil trajectories of the anterior (blue) and posterior (red) centrosomes after laser ablation for different species were averaged (filled circles). Error bars: SEM. The mean data was fit to the Kelvin-Voigt model (bold line), Equation 1. The faint lines represent individual profiles of anterior (blue) and posterior (red) centrosomes. **(C)** The parameters force F (left) and elasticity k (right) obtained from the KV-model fit are plotted as box-plots. White circles: mean value for a species, red/blue circles: estimates from fits to individual recoil trajectories. The KS test was applied to inter-species and A/P comparisons for significance. The asterisks indicate * p<0.05, ** p<0.01, *** p<0.001, **** p<0.0001 and ns: not significant.

where F is the effective force driving the movement of the centrosome as it recoils after cutting, k is the elasticity of the medium and g is the drag coefficient. Assuming the centrosome and MT aster can be treated as a spherical object, the Stokes drag coefficient can be estimated from the expression:

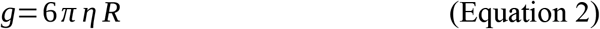

here η is the viscosity of the medium and R represents the centrosome aster radius.

In the Kelvin-Voigt model, the centrosome position is governed by a pulling force F, here exerted on astral microtubules by cortical force generators and resisted by the elasticity k that results from a combined effect of cytoplasmic components, actin meshwork and astral MTs and the viscous drag *g*, that measures the opposition to the motion of the aster. Given the free parameters, a good fit to the recoil trajectories requires constraining the fit. We use our measurement of the effective cytoplasmic viscosity to constrain the drag coefficient *g* (Equation 2).We also measured centrosome sizes from images based on DIC microscopy, where centrosomes appear as a smooth disk that excludes the cytoplasmic granules due to the high density of astral microtubules around the pericentriolar material. The radius of this disk is taken as the centrosome size. Variations in centrosome radius were very limited between species, ranging from 2 to 3.59 μm (Table S1). Implicit in our approach is the assumption that centrosome size remains constant during anaphase. Substituting these values into Equation 2, we reduced the free parameters of the fit to only two: the force F and rigidity k. We proceeded to fit the individual data from multiple experiments and develop an understanding for the diversification of spindle mechanics.

Our results revealed the net pulling force acting on the anterior and posterior centrosome of *C. elegans* spindles to be 45.8 pN and 49 pN, respectively (Fig. 4C, Table 2). We found *C. elegans, C. remanei* and *C. monodelphis* spindles experience comparable magnitudes of forces. However, compared to C. elegans, those in *P. pacificus* and *O. tipulae* are lower, while *D. sp*.*1* appears to have the lowest values measured amongst all 6 species. Except for this 10 fold difference between *C. elegans* and *D. sp. 1*, changes in pulling forces are however very limited for the other species, with a fold change of ∼2.

**Table 2.** Average of parameters obtained by fitting the individual trajectories of centrosomes after spindle cut to the Kelvin-Voigt model seen in Fig. 3B.

| Table 2 |  |  |  |  |
| --- | --- | --- | --- | --- |
| *Species* | *Force (pN)* |  | *k (pN/μm)* |  |
|  | *Anterior* | *Posterior* | *Anterior* | *Posterior* |
| *C. elegans* (n = 7) | 49.99 ± 21.06 | 53.49 ± 22.94 | 8.31 ± 2.50 | 9.81 ± 3.03 |
| *C. remanei* (n = 6) | 32.32 ± 12.42 | 30.29 ± 13.56 | 5.20 ± 1.72 | 6.35 ± 2.74 |
| *C. monodelphis* (n = 14) | 63.60 ± 26.60 | 49.87 ± 25.50 | 10.66 ± 5.58 | 9.76 ± 8.05 |
| *D. sp. 1. JU359* (n = 3) | 4.84 ± 2.92 | 9.31 ± 7.90 | 4.87 ± 6.32 | 5.27 ± 6.12 |
| *P. pacificus* (n = 13) | 28.13 ± 13.25 | 33.85 ± 19.93 | 4.76 ± 3.95 | 5.98 ± 4.41 |
| *O. tipulae* (n = 11) | 29.13 ± 13.76 | 32.88 ± 13.18 | 7.70 ± 4.84 | 5.24 ± 2.96 |

The rigidity (k) values range between 4 to 10 pN/μm which is consistent with values previously obtained for *C. elegans* embryos at the stage of spindle centering (Garzon-Coral et al. 2016). Although variations of the rigidity appear even more constrained than pulling forces, they show a similar trend between species. Here, only *P. pacificus* and *O. tipulae* show clear differences with C. elegans (Fig. 4C, Table 2).

The mechanical differences in spindles between species in terms of pulling forces, elasticity and viscous drag may suggest a global trend that might explain differences in unperturbed spindle behavior. To investigate this we proceed to correlate these variables in order to find patterns.

### Towards the definition of a parameter space for spindle positioning and oscillations

Overall, our results show substantial and independent variation from one another, even between closely related species (Fig. 3E and 4C, Tables 1 and 2). Interestingly we find the spindle recoil dynamics measured by velocity, v (Fig. 5A) and time-constant τ after cutting (Fig. 5B) correlate strongly with effective cytoplasmic viscosity (|r| > 0.8). To our surprise even cell cycle time shows a similar correlation with viscosity with r > 0.8 (Fig. 5C, see Discussion). At the same time, there is poor correlation between viscosity and estimated spindle pulling force (Fig. 5D) and elasticity (Fig. 5E). The recoil velocity and time are poorly correlated with cell length.

**Figure 5.**
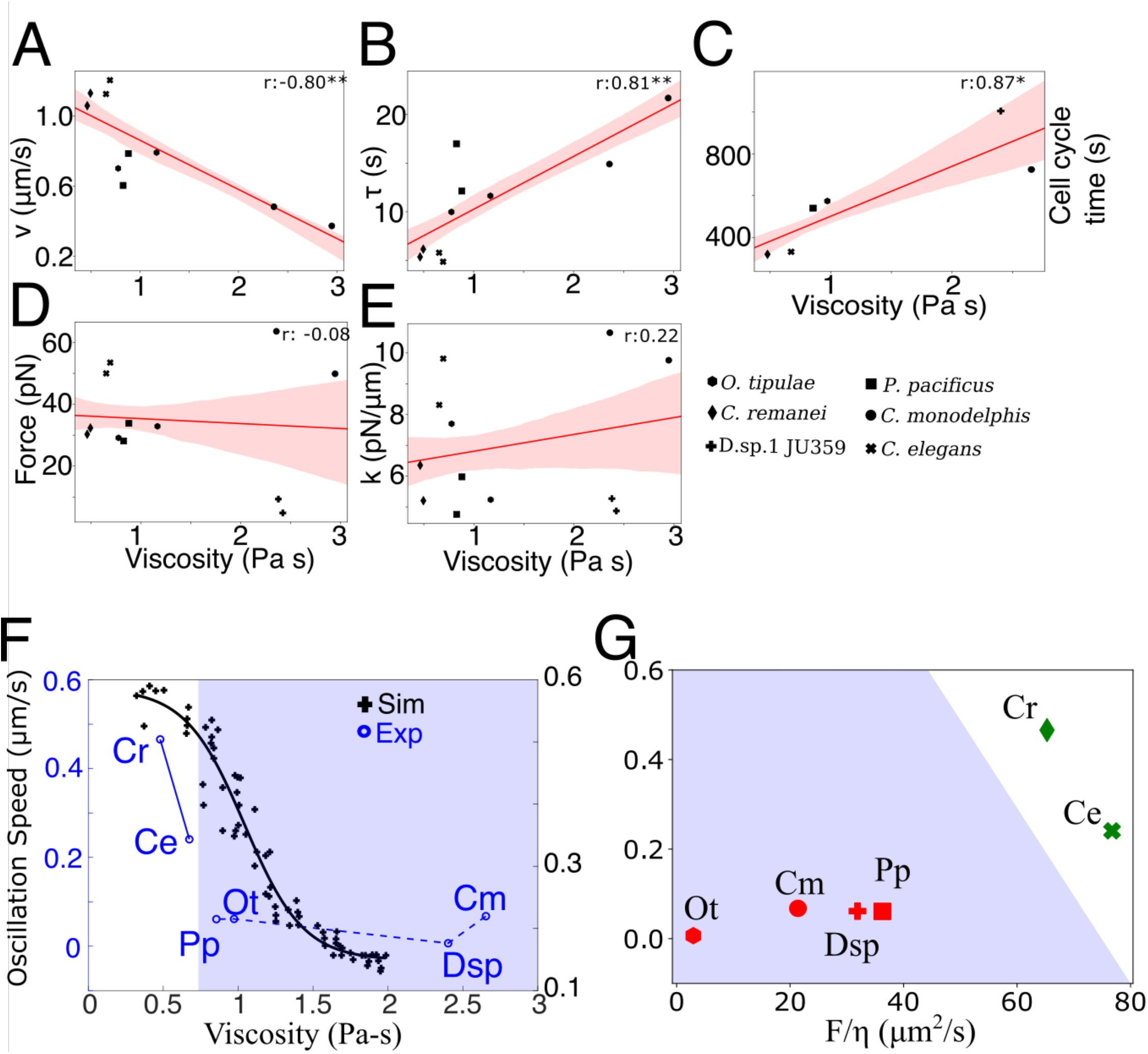
Effect on spindle mobility of divergence in cell mechanics. The covariation in all 6 species of effective viscosity with **(A)** recoil velocity v, **(B)** decay constant τ, **(C)** cell cycle time and **(D)** spindle pulling force (pN) and **(E)** elasticity (pN/μm) of the half-spindle are plotted. Symbol represent individual species. The red line indicates regression fit with one standard deviation (confidence interval 0.68) indicated by the shaded area. r indicates Pearson’s correlation coefficient. Asterisks indicate significance level * p<0.05, ** p<0.01, *** p<0.001, **** p<0.0001. **(F)** Spindle oscillation speeds of embryos of each species (described in Figure S4) were plotted as a function of the effective viscosity (circles, Ce: *C. elegans*, Cr: *C. remanei*, Ot: *O. tipulae*, Pp: *P. pacificus*, Dsp: *Diploscatper sp. 1* and Cm: *C. monodelphis*) and compared to simulation predictions (+) from previous work (Kozlowski et al., 2007). The simulation data is fit to a four parameter sigmoid function (Equation 9) with fit parameters a= 0.44 μm/s, d= 0.15 μm/s, c= 1.04 Pa-s and s= 0.2 Pa-s. **(G)** The spindle oscillation speeds are also plotted as a function of the ratio of the spindle pulling force and the viscosity. The shaded region in **(F)** and **(G)** represents those species whose spindles do oscillate.

In the crowded environment of the cell, it is not surprising that viscosity plays an important role in intracellular mobility. Therefore, when we measure the unperturbed spindle oscillation speed (Fig. S5) and correlate it with viscosity, we find a distinct separation on the basis of viscosity between species that show bonafide oscillations (Ce, Cr) and the rest (Pp, Ot, Dsp and Cm) which occurs above a viscosity of ∼0.6 Pa s (Fig. 5F). This trend is surprisingly comparable to a computational model (Kozlowski et al., 2007) which had predicted a steep decrease in oscillation speed when viscosity increased above ∼0.6 Pa s. The predictions in previous work had never been tested due to the difficulty of such experiments and the general nature of viscosity. Our results show this is possible to challenge the model with measures obtained with comparative biophysics.

The lack of a complete fit between the predictions and data (Fig. 5F) suggests potentially many unknowns in the model and possibly a missing variable. Since spindle movement is driven by the pulling forces, a tradeoff between viscosity and forces might be necessary to sustain oscillations. This is supported by the case of *C. remanei* in which spindles oscillate whereas both forces and viscosity are reduced compared to C. elegans (Fig. 3E, 4C). On the other hand, *P. pacificus* experiences lower forces compared to *C. elegans* but identical viscosity which could explain the absence of oscillations. In order to account for a potential tradeoff between the pulling forces and viscous drag that opposes it, we estimate the ratio between the two (ω), to examine if it may predict the propensity of spindles to oscillate.

We define it as:

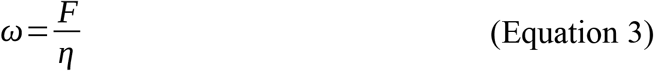

where F is the spindle pulling force and η is the effective viscosity. We find spindle oscillation speed plotted as a function of this ratio shows a clear separation between the species that correlates with spindle oscillations (Fig. 5G). A low value of the ratio is suggestive of either low force or high viscosity, while a high value indicates either low viscosity or high force. This segregation of species in terms of the relative tradeoff in viscosity and force, appears to predict the oscillations of anaphase spindles in the species examined.

## Discussion

The intracellular aqueous environment of macromolecules and presence of cytoskeletal proteins has made the study of viscous and elastic properties of cells vital for our understanding of cellular mechanobiology. However, the diversity in biophysical properties of cells, over the course of evolution, is still unclear. In this study we aim to explore the reason why spindle motion varies between closely related species of nematodes. We had previously identified species for which spindle elongation and displacement at anaphase is accompanied by transverse oscillatory movements, similar to the situation found in *C. elegans*. In contrast, many nematode species do not undergo these oscillations, despite an identical off-center displacement of the spindle at anaphase. Comparing spindle movements is a difficult task because spindle elongation, displacement and oscillations occur simultaneously. Moreover, the spindle is composed of two centrosomes that are oscillating in an anti-phase manner and which are linked by a central spindle, whose mechanical properties are mainly unknown. Based on previous work demonstrating the role of motor-driven pulling forces (elastic) and the effect of cytoplasmic drag in damping this movement (viscous), we proceeded to examine the comparative viscoelastic properties of the spindle motion of multiple species. As a first step, we laser-ablated the central spindle and analyzed the recoil trajectories to estimate velocity and characteristic time constants. However, a viscoelastic model required an estimate of viscosity between species. Using the diffusive motion of high-contrast cytoplasmic granules, we found a ten-fold variation between the viscous drag. Using the drag to constraint a viscoelastic model of spindle recoil, we find spindle elasticity to be largely conserved but pulling forces to vary in upto ten-fold between species. Correlation of intact spindles as well as recoil dynamics suggests a balance of forces acting on spindles determines the mobility. Cellular parameters such as centrosome size or cell length do not appear to correlate with spindle retraction parameters, while viscosity strongly correlates with both recoil parameters and intact spindle mobility. However, a complete picture is only obtained when we consider the relative proportion of pulling forces and viscous drag. This ratio appears to determine the emergence of spindle oscillations above a critical value, that could serve as a prediction of biophysical constraints on the evolution spindle oscillations.

Our results first demonstrate that regardless of the presence of spindle transverse oscillations, all studied species are subjected to cortical pulling forces. Interestingly, although dynein, is a highly conserved protein, the protein responsible for its anchoring at the cortex, i.e. GPR-½ or LIN-5, are not found in the genomes of *P. pacificus, O. tipulae* or other members of the *Diploscapter* genus (Delattre and Goehring, 2021). This raises the possibility that although the pulling machinery is conserved, the molecular complex responsible for this force has changed between closely related species. We found variations in the net pulling forces, by a factor of ∼10, between species. Previous models have proposed that modulation of the force can be achieved by changes in the number of motors, the individual force per motor, the attachment and detachment rate of motors, and microtubule dynamics, as seen in other systems (Sutradhar et al., 2015, Jain et al., 2021). Establishing transgenic lines, in particular to follow live microtubules, in these different species is a necessary step towards more quantitative measurements of these parameters.

We also find the rigidity (spring elasticity) parameter differences between species vary only over a factor of two, ranging between 4.8 and 9.8 pN/μm. In terms of the cell, this parameter can be understood to be a combined measure of cytoplasmic stiffness and microtubule components. This lack of clear difference suggests that the density of microtubules and actin meshwork are likely to be comparable between closely related nematode species in the same cell-cycle stage. A fluorescence based approach to compare these mesh-works could test this hypothesis in future. This result suggests that the pulling forces and drag are the primary determinants of the spindle behavior in these species.

Finally, we found very large differences in cytoplasmic viscosity between species using a non-invasive method. Early work measuring cytoplasmic viscosity in vertebrate cell lines reported a viscosity of 0.282 Pa.s following the Brownian motion of cytoplasmic inclusions (Alexander and Rieder, 1991). A more recent study using micron-sized beads and magnetic tweezers to measure the viscoelastic properties of the cytoplasm in *C. elegans* during spindle positioning also reported viscosity of 0.159 Pa.s (Garzon-Coral et al., 2016). However, tracking microrheology of the mobility of injected nanospheres in one-celled embryos of *C. elegans* reported a spatially uniform cytoplasmic viscosity of 1 Pa.s (Daniels et al., 2006). Accumulating evidence of the probe size dependence of such measurements due to macromolecular crowding (Etoc et al., 2018; Mogilner and Manhart, 2018) could explain this order of magnitude difference reported by different workers, in the same species. Our measurements retrieve a viscosity value that is in between these two extremes for *C. elegans*. We report the effective cytoplasmic viscosity to be ∼600 times higher than the viscosity of water. Using the same approach to multiple species, we have uncovered large variations, up to an order of magnitude, between species with *C. monodelphis* showing the highest viscosity. The differences we measure could arise from multiple factors such as differences in protein concentrations, presence of different densities of cytoskeletal meshwork or higher organelle densities. In future a careful morphological comparison between the species could potentially allow us to address the question of how these differences in viscosity could arise.

We also note that species that have high viscosity also have a longer cell cycle duration (Fig. 5C, correlation coefficient R= 0.87). This result suggests that the only combinations that have been retained by natural selection are compensatory changes, where high viscosity is compensated by a slow down of the cell cycle or vice-versa. Over the course of evolution, slow species could have afforded an increase in cytoplasmic viscosity because even though objects are slowed down by the viscous drag, they will have time to reach their final position. Conversely, a low viscous drag may have preconditioned the emergence of a fast cell cycle. Regardless of the orientation of changes, this interesting correlation raises the question of the selective pressure responsible for species-specific viscosity values.

Overall, we found variations in all parameters and species-specific combinations of parameters, which are all compatible with asymmetric spindle positioning. How far these parameters can change without perturbing the first embryonic division remains an open question but our study is a first step towards the exploration of this parameter space. Already with a small number of species, we found that viscosity and pulling forces can change by an order of magnitude. The specific case of *D. species 1*, for which all parameters have changed dramatically compared to *C. elegans*, demonstrate how much changes can be tolerated without affecting asymmetric cell division.

Back to our initial question, our results allow us to define which combination of parameters are now compatible with spindle transverse oscillations. Reduction of pulling forces can still lead to oscillations provided it is compensated by reduced cytoplasmic viscosity, as seen in *C. remanei*. However, changes in a single parameter, as seen in *P. pacificus* (reduced pulling forces) or in *C. monodelphis* (higher viscosity), critical for oscillations, do not suffice. We propose that a tradeoff between cortical pulling forces and cytoplasmic viscosity results in spindle oscillations, when the ratio of pulling forces to viscosity are high. This provides a framework that could both be tested with more species of nematodes, as well as generalized to other cellular systems.

## Materials and methods

### Image acquisition of nematode embryos and strain maintenance

All strains were maintained at 20°C on Nematode Growth Media seeded with *E. coli* OP50, as described in (ref Brenner): *Caenorhabditis elegans* (N2), *Caenorhabditis remanei* (PB219), *Caenorhabditis monodelphis* (SB341), *Diploscapter sp. 1* (JU359), *Oscheius tipulae* (CEW1), *Pristionchus pacificus* (PS312). (Valfort et al., 2018). For embryo recording, females were dissected in M9 and one-cell stage embryos were placed between slide and cover slip on a 2% agar pad (ref). Embryos were observed with a Zeiss Axioimager A1 or A2 with a 100X DIC Plan Apochromat NA 1.4 lens. For video recording of the cell division, we took 2 images per second. with a digital Kappa camera (DX4-285FW).

### RNAi experiments

RNAi experiments on *C. elegans* were performed by feeding, as described in (ref). Wild type L4 larvae were fed for 24 hours, with HT115 bacteria producing the double-strand RNA of atp-2 and cyc-1 genes. We considered that RNA interference was achieved when the one-cell embryos did not show signs of spindle movements during mitosis.

### Laser ablation

One-cell embryos in prophase or prometaphase were mounted between slide and coverslip as described above. Embryos were then recorded on an inverted spinning disk confocal microscope (Leica DMI4000B-CSU 22 Yokogawa) with a 100X immersion objective (HCX PL APO 1.4 oil) controlled by Metamorph. Images were acquired with an iXon3 897 Andor camera every 0.5 seconds. Spindle severing was performed using a UV laser module (lambda=355 nm) iLas2 Roper, as described in (Grill 2001). For each species, the laser power was adjusted so that the cut, performed at the onset of spindle elongation, generated a rapid movement of the centrosomes (due to spindle severing), but did not arrest the cells (due to excess laser power).

### Granule-based viscosity measurement

DIC images of nematode zygotes were acquired every 0.5 s in the mid-plane of the embryo during prometaphase. Images were pre-processed to enhance contrast (with 0.3% saturated pixels) using Fiji (Schneider et al., 2012). Granule motion was analyzed by tracking whole embryos and cropped regions of interest (ROIs) using a home-built program in MATLAB (Mathworks, USA) for single particle segmentation and tracking of DIC images (Chaphalkar et al., 2021). Granule data was averaged with ∼2000 granules per embryo with between 5 and 19 embryos per species analyzed. The MSD of particles was calculated using the x-y coordinates of tracked granules as follows:

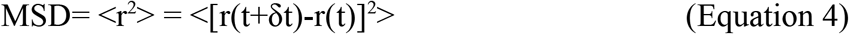

Here, *r* is the displacement of the particle at two time-points separated by a time-step *δt*. We employ a sliding window approach and estimate the MSD for the first 3/4th of the data to avoid artefacts due to undersampling at large values of *δt* (Michalet, 2010; Khetan and Athale, 2016). The effective diffusion coefficient (*D*) was estimated by fitting the linear region, up to 5 seconds of the average MSD profile to the anomalous diffusion model:

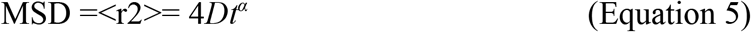

Here, *t* is the time-step and *α* is the anomaly parameter that indicates the nature of diffusion. The motion is said to be purely diffusive if α ≈ 1, sub-diffusive or ‘restricted’ when α < 1 and super-diffusive or ‘transported-like’ when α > 1. The fluid viscosity (ηf) was estimated from the effective diffusion coefficient (*D*) and radius (r) of granules using the Stokes-Einstein relation:

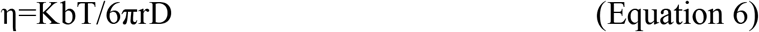

Granule radii (r) are very similar ∼0.2-0.3 μm across species (Fig. S3) and are used to estimate the viscosity for each species. In order to account for the crowded nature of the cytoplasm due to granule packing the effective viscosity (η_eff_) was estimated based on the approximation for soft-spheres (Quemada, 1977) by correcting for the packing fraction of the embryo due to the 2D granule packing fraction (ϕ). The granule packing fraction for each species was calculated from 8 representative ROIs each from the anterior and posterior regions, with ∼400 granules per species. The granule density per unit area, ϱ_g_=N_g_/A_cell_, was estimated for anterior and posterior 1/3rd of each embryo along the major axis in each species (Fig. 2D). The area of each granule (Ag) was measured from the granule radius to arrive at the granule packing fraction ϕ_2D_ = ϱg*Ag and used to calculate the effective viscosity as follows:

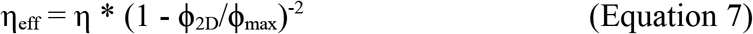

where ϕ_max_ is the maximal packing fraction, taken to be 0.64 for random packing (Buscall et al., 1994) and η is the solvent phase viscosity. The packing fraction was measured from DIC images in the anterior and posterior regions of each species (Fig. 2D).

### Oscillation speed and frequency

Spindle oscillations were analyzed in representative trajectories of each species by smoothing the position from centerline with time using the discrete cosine transform to reduce low-frequency noise with the threshold greater than 1.5 for *C. monodelphis* and *O. tipulae*, and 0.56 for the remaining species. The speed of oscillation was calculated as the change of position over successive windows of 5 second intervals, throughout the trajectory. Oscillation frequency was estimated using the FFT on the smoothed data from all species. Mean and max oscillation speed was calculated at a 5 second interval over the smoothed trajectories.

### Data fitting

The initial recoil velocity and rate of decay form the recoil trajectories was obtained by fitting the anterior and posterior centrosome recoil trajectories to the function:

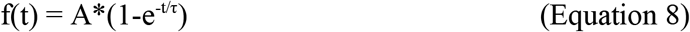

where A is amplitude of recoil and τ is decay constant of the exponential and V is the recoil velocity given by *V = A/τ* based on previous work (Sumi et al., 2018). Oscillation speed predictions from simulations as a function of increasing viscosity were taken from previous work by Kozlowski et al., (2007) by digitizing the plot (webplotDigitizer) and fitting it to a 4-parameter sigmoid function described previously (Khetan and Athale, 2016):

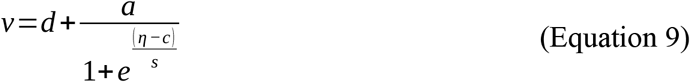

Where v is the speed, η is the viscosity, d and a are the minimal and maximal speeds, c is the half-maximal viscosity and s is the steepness of the profile. Individual data from centrosome recoil of all species were ignored if the goodness of fit measure (R^2^) was less than 0.8, except for *D*.*sp*. where this cutoff was 0.5. As a result of this we do not report v and τ values for this species (Fig. 5A, B).

### Statistical tests and correlation

All correlations were performed using Pearson correlation test. *Regplot* was used to illustrate regression fits for all correlations (Seaborn 0.11.0, Python3). The size of the confidence interval was set to 68 which falls within 1 standard deviation (Fig. 5A-E). All statistical tests were performed in SciPY 1.5.2. Correlations were quantified using the function for Pearson’s correlation coefficient *pearsonr* in Python (SciPy). Viscosity between species and within a species across A/P regions were compared using the KS test.

## Acknowledgements

Anushree Chaphalkar and Tanmaya Sethi were involved in the early stages of the project. CA is grateful to Vijaykumar Chikaddi for discussions. MD thanks Fabien Montel for critical comments and discussions. MD and TB acknowledge the contribution of the imaging platform PLATIM from SFR Biosciences Lyon.

## Supporting Tables

**Table S1.** The cell length (long axis) in μm, cell cycle duration in seconds and centrosome radius of the anterior and posterior centrosomes is reported for the 6 species analyzed in this study.

| Species | Cell length<br>($\mu\text{m}$) | Cell cycle<br>duration (s) | Centrosome Size ($\mu\text{m}$) | |
| --- | --- | --- | --- | --- |
|  |  |  | Anterior | Posterior |
| *C. elegans* | 50.59 | 331 | 3.6 | 3.26 |
| *C. remanei* | 48.60 | 319 | 3.06 | 2.95 |
| *C. monodelphis* | 50.63 | 725 | 3.25 | 2.88 |
| *D. coronatus* | 38.15 | 1005 | 2.23 | 2.05 |
| *P. pacificus* | 54.04 | 540 | 3.19 | 2.96 |
| *O. tipulae* | 46.98 | 574 | 2.83 | 2.77 |

**Table S2.** The spindle cutting experiments tracks were fit to a model of recoil kinetics (Equation 8) to obtain the recoil velocity and time-constant (mean ± SEM) for each species, for anterior and posterior centrosomes (details in the Methods section).

| Species | Retraction velocity (um/s)<br>(Mean ± SEM) | | $\tau$ (s)<br>(Mean ± SEM) | |
| --- | --- | --- | --- | --- |
|  | Anterior | Posterior | Anterior | Posterior |
| *C. elegans* | 1.12 ± 0.12 | 1.23 ± 0.17 | 6.16± 0.70 | 4.59 ± 0.52 |
| *C. remanei* | 1.13 ± 0.09 | 1.06 ± 0.2 | 5.88 ± 0.83 | 5.37 ± 1.02 |
| *C. monodelphis* | 0.46 ± 0.05 | 0.38 ± 0.05 | 14.94 ± 1.84 | 20.98 ± 4.87 |
| *D.sp.1* | 0.05 ± 0.03 | 0.074 ± 0.04 | 23.77 ± 15.28 | 23.09 ± 5.50 |
| *P. pacificus* | 0.59 ± 0.07 | 0.73 ± 0.1 | 15.42 ± 3.49 | 10.91 ± 2.39 |
| *O.tipulae* | 0.70 ± 0.1 | 0.75 ± 0.1 | 9.98 ± 3.02 | 15.27 ± 4.54 |

## Supporting Figures

**Figure S1.**
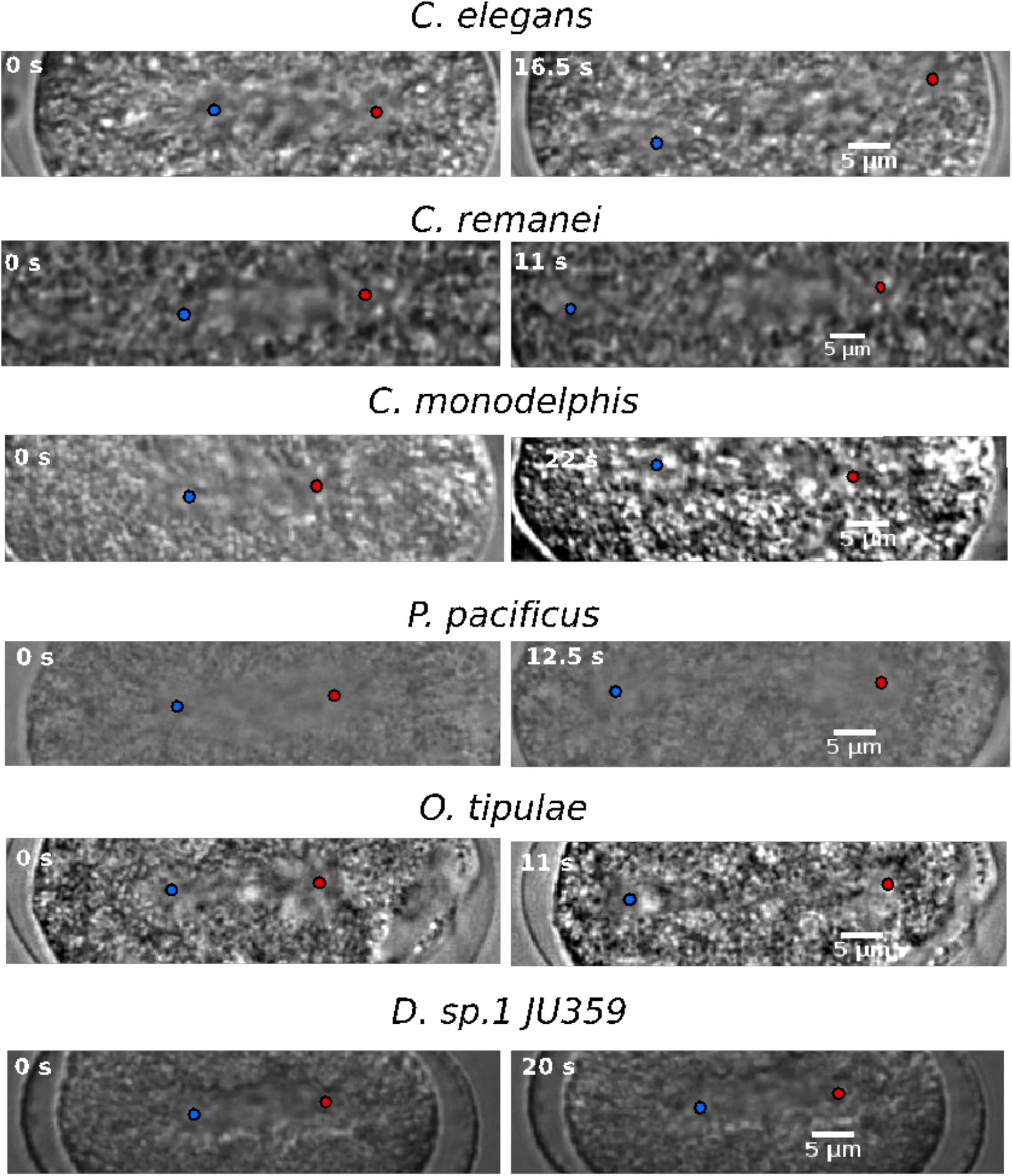
Effect of spindle cutting on centrosome recoil. Representative images from immediately before (left column : t = 0 s) and after spindle cutting (right column) for different species are shown. The images after cutting are selected based on the approximate time required for the centrosomes to achieve their maximal displacement from the initial position. Scale bar 5 μm.

**Figure S2.**
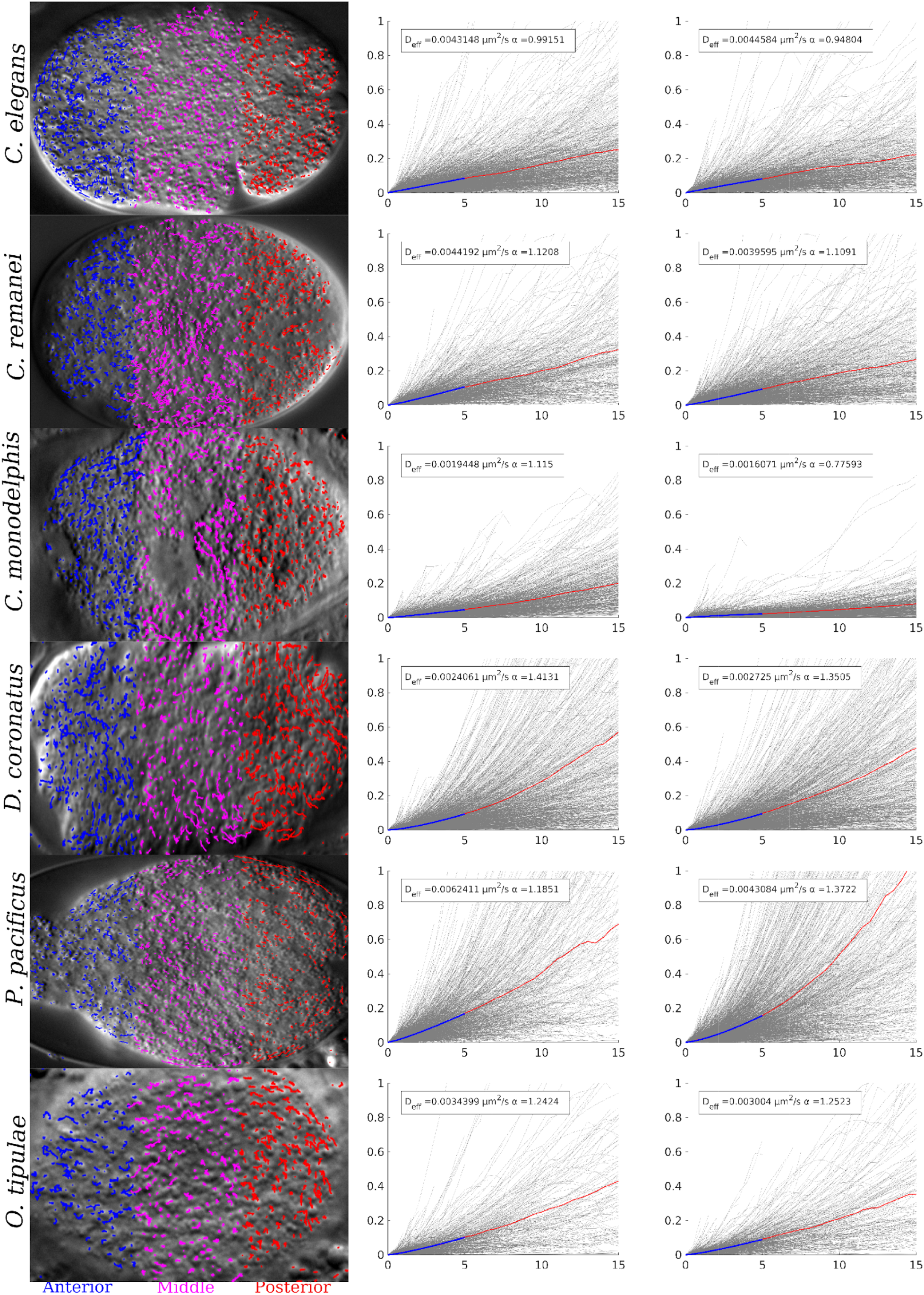
Measuring granule diffusion coefficient for multiple species. Montage of tracked granules from individual species and corresponding MSD analysis for anterior (left) and posterior (right) regions. Anterior (Blue), middle(Pink) and posterior (Red) regions correspond to 33% of the major axis length. Insert values show Deff and α values from the anomalous diffusion fit (till 5s) to the mean MSD curve (red). Individual frames have been resized to fit and are not to scale.

**Figure S3.**
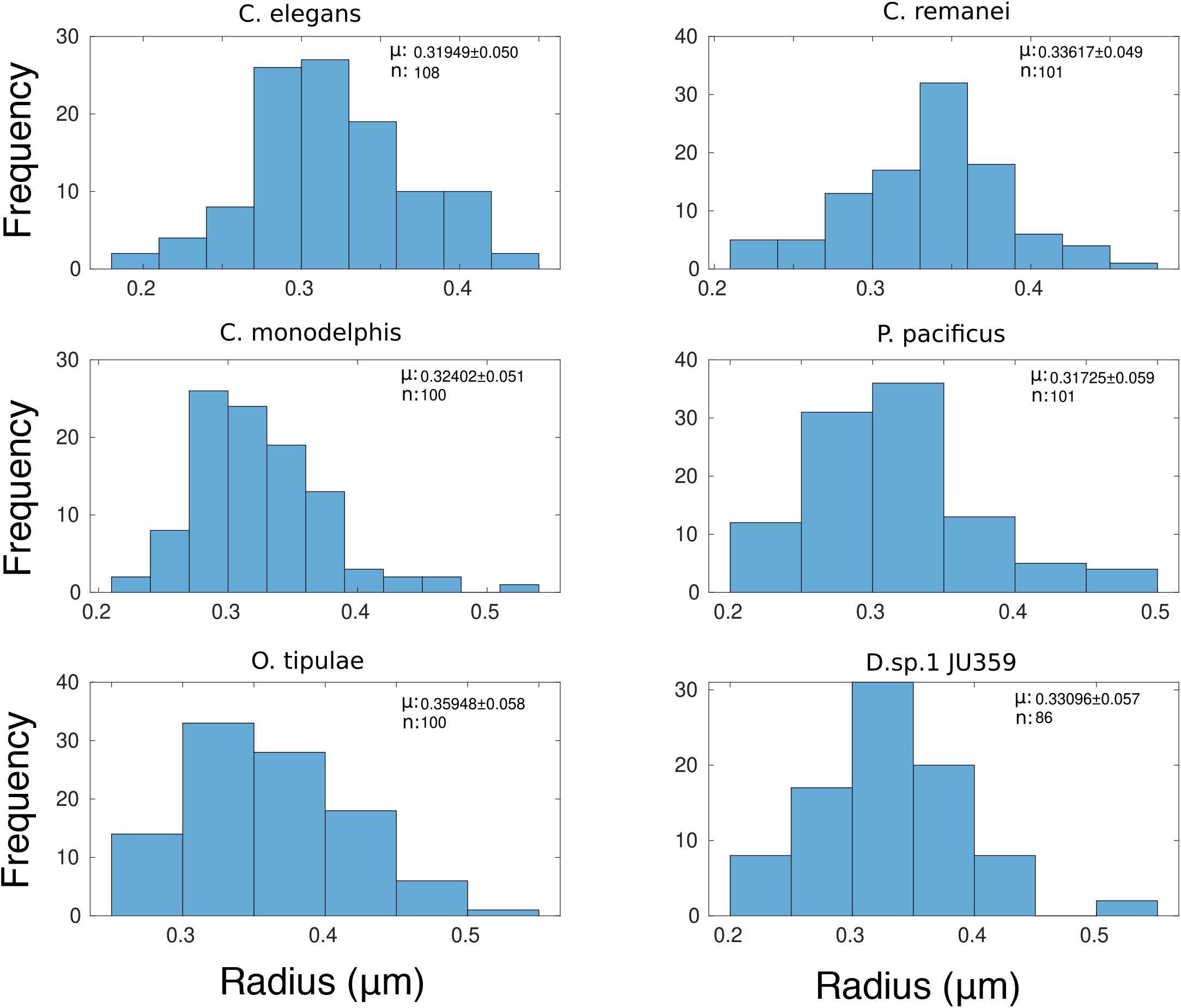
Differences between granule radii of different species. The granule radii of multiple species were measured interactively by averaging granule sizes across ROIs and multiple individual embryos (n ≈ 100) for each species.

**Figure S4.**
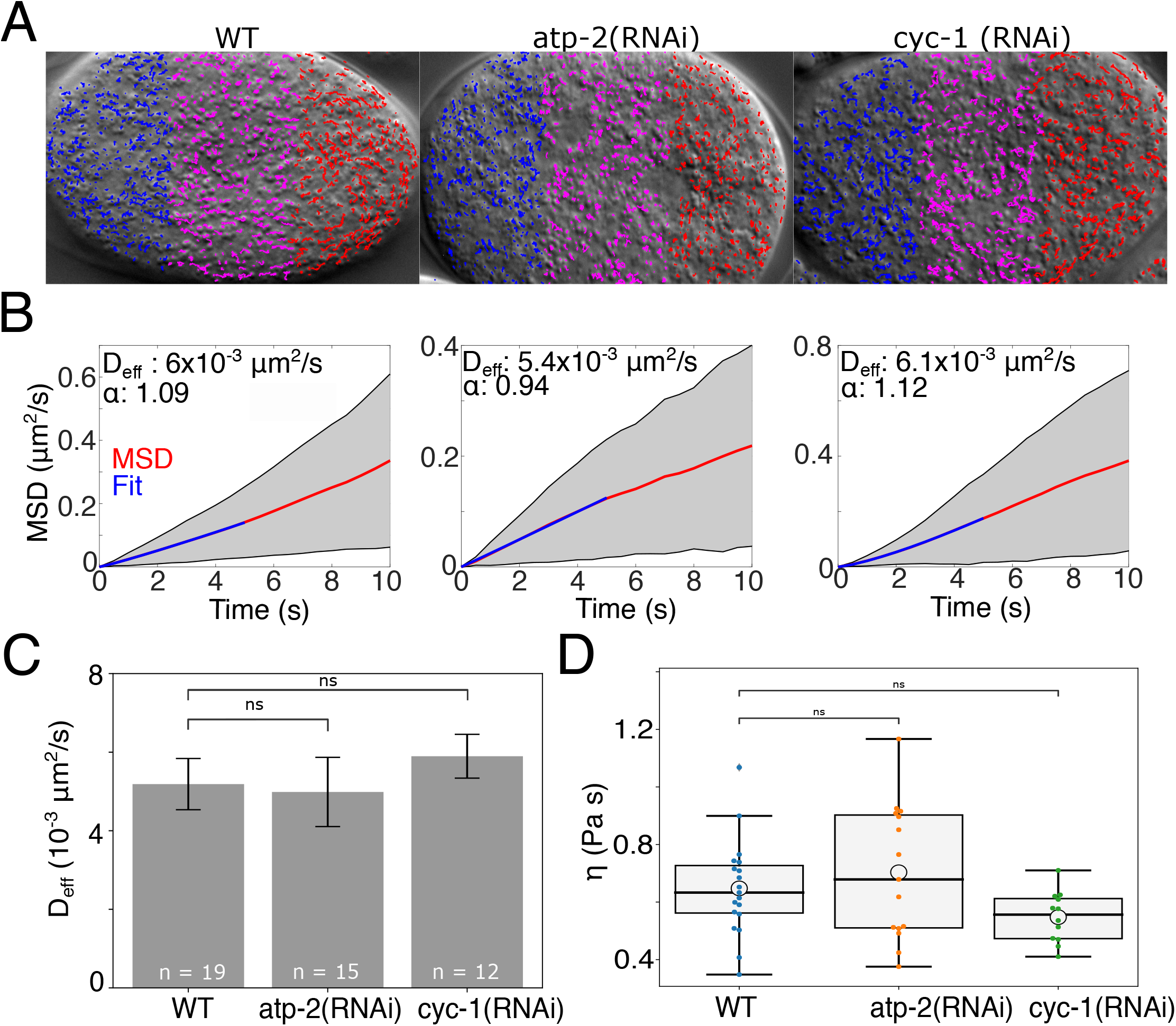
Effect of ATP depletion on granule mobility. **(A)** Representative tracked granules from WT are compared to embryos treated with RNAi targeting ATP2 and CYC-1 are overlaid on DIC images of the embryo in the anterior (blue), mid-cell (magenta) and posterior (red) regions. **(B)** The MSD (red, Equation 4) with time is plotted for whole embryos corresponding to **(A)**. The grey area is the s.d. The data was fit using the anomalous diffusion model, Equation 5 (blue) to estimate the effective diffusion coefficient (D_eff_) and anomaly coefficient (α). **(C)** The mean D_eff_ (error bar: 2 SEM) and **(D)** effective viscosity are compared between untreated and treated embryos. The mean values were compared using a KS test; ns: not significant.

**Figure S5.**
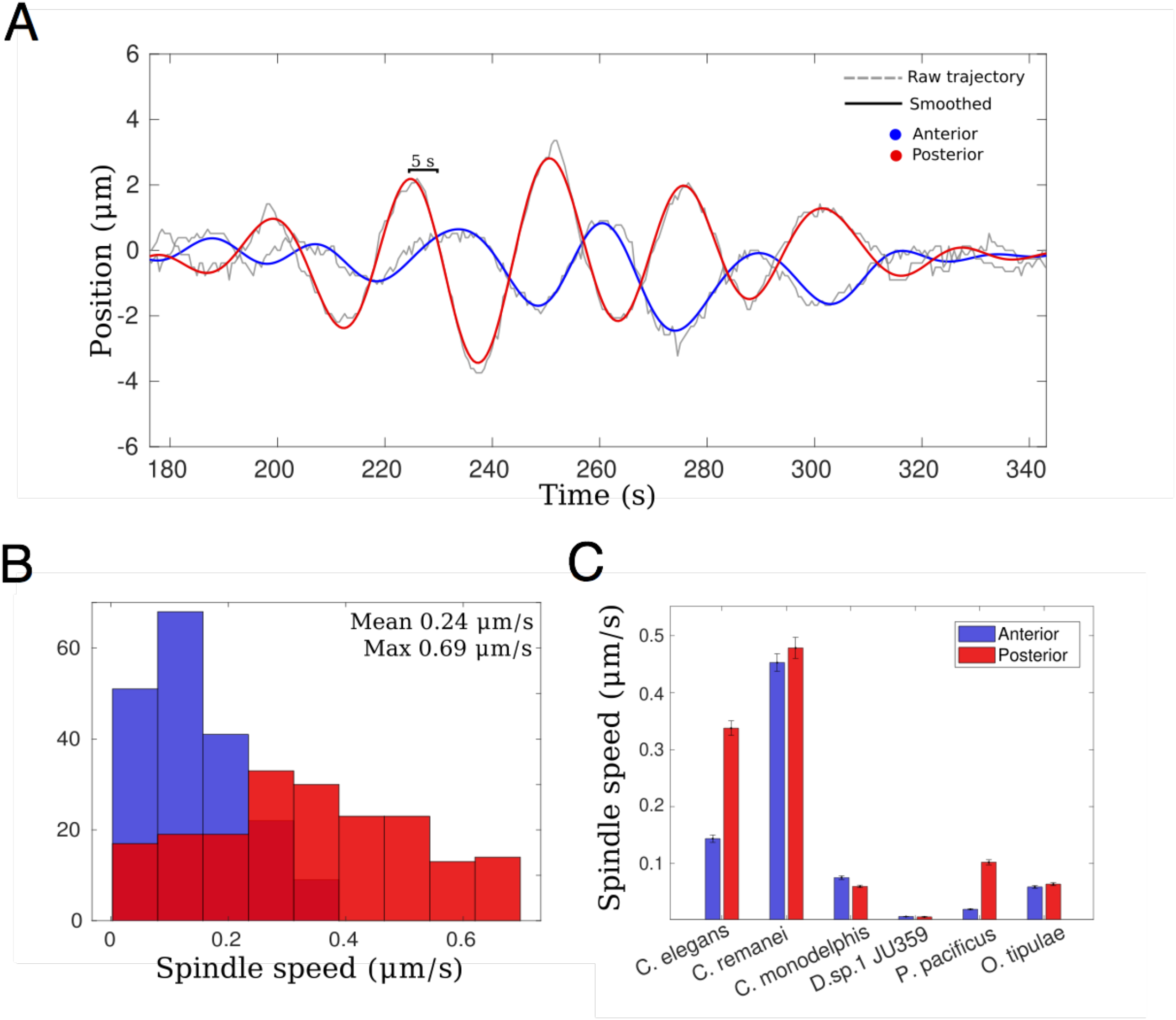
Spindle oscillation speed. **(A)** The spindle pole positions of a representative *C. elegans* embryo are plotted relative to the centerline as a function of time (black line) for anterior and posterior poles. Small fluctuations are removed by smoothing the data (blue: anterior, red: posterior). Oscillation speed is estimated as a change in position in a 5 s window successively throughout the trajectory from the smoothed trajectory. (B) The frequency distribution of anterior (blue) and posterior (red) centrosomes is plotted. (C) Speeds (mean ±SEM) are compared between the 6 species.

## Supporting Videos

**Video S1(A-F)**. Tracked granules in a DIC time series of studied embryos during first mitotic division are shown. Trajectories are color coded based on 1/3 of the major axis length into anterior (blue), posterior (red) and middle (magenta) region. Species are in order **(A)** *C. elegans*, **(B)** *C. remanei*, **(C)** *C. monodelphis*, **(D)** *D*.*sp*.*1 JU359*, **(E)** *P. pacificus* and **(F)** *O. tipulae*. Scale: 5 μm; Δt: 0.5 s.

**Video S2. (A-C)** DIC time series of the RNAi treated *C. elegans* embryos to deplete **(A)** atp-2 and **(B)** cyc-1 compared to **(C)** untreated (wild type), were tracked to follow granule mobility. Granule trajectories are overlaid with colors indicating location along the major axis. The anterior (blue), posterior (red) and middle (magenta) regions were determined as 1/3rd of the length of the AP axis of the embryo. Scale: 5 μm; Δt: 0.5 s.

